# Loss of CD109 Amplifies NF-κB Signaling and Inflammatory Reprogramming in Dermal Fibroblasts

**DOI:** 10.64898/2026.07.03.736423

**Authors:** Adel Batal, Saniya Pamnani, Shufeng Zhou, George Bou-Gharios, Anie Philip

## Abstract

Fibroproliferative diseases such as systemic sclerosis are complex conditions characterized by chronic skin inflammation and progressive fibrosis, with fibroblast activation as a central feature. While Transforming Growth Factor Beta (TGF-β) signaling is a well-established driver of fibrosis in SSc, inflammatory pathways such as Nuclear Factor Kappa B (NF-κB) also contribute substantially to disease morbidity. We previously identified CD109 as a TGF-β co-receptor and negative regulator of fibrotic signaling; however, its role in inflammatory signaling remains unknown. Here, we investigate the function of CD109 in regulating inflammatory signaling in skin fibroblasts. We show that, CD109 co-localizes and associates with Toll-like receptors (TLR2, TLR4) and tumor necrosis factor receptors (TNFRI, TNFRII), and that loss of CD109 enhances TNF-α–induced NF-κB activation and reprograms cytokine production in human dermal fibroblasts. Furthermore, both global and fibroblast-specific CD109 knockout mice exhibit increased immune cell infiltration and skin inflammation. In parallel, single-cell transcriptomic analyses across a pan-disease fibroblast atlas show that CD109 expression is preferentially maintained in structural and homeostatic fibroblast subtypes, whereas immune-interacting fibroblast subsets consistently display decreased CD109 levels. Pathway-level analyses of fibroblast pseudobulk samples reveal altered activity of canonical inflammatory pathways in SSc compared to healthy skin. Together, these findings identify CD109 as a fibroblast-intrinsic negative regulator of inflammatory signaling and suggest a broader role for CD109 in modulating inflammatory responses in systemic sclerosis.

**Graphical Abstract:** CD109 Restrains Fibroblast-Driven Inflammation by Modulating NF-κB Signaling. Generated using *FigureLabs.ai* and edited using Adobe Photoshop.

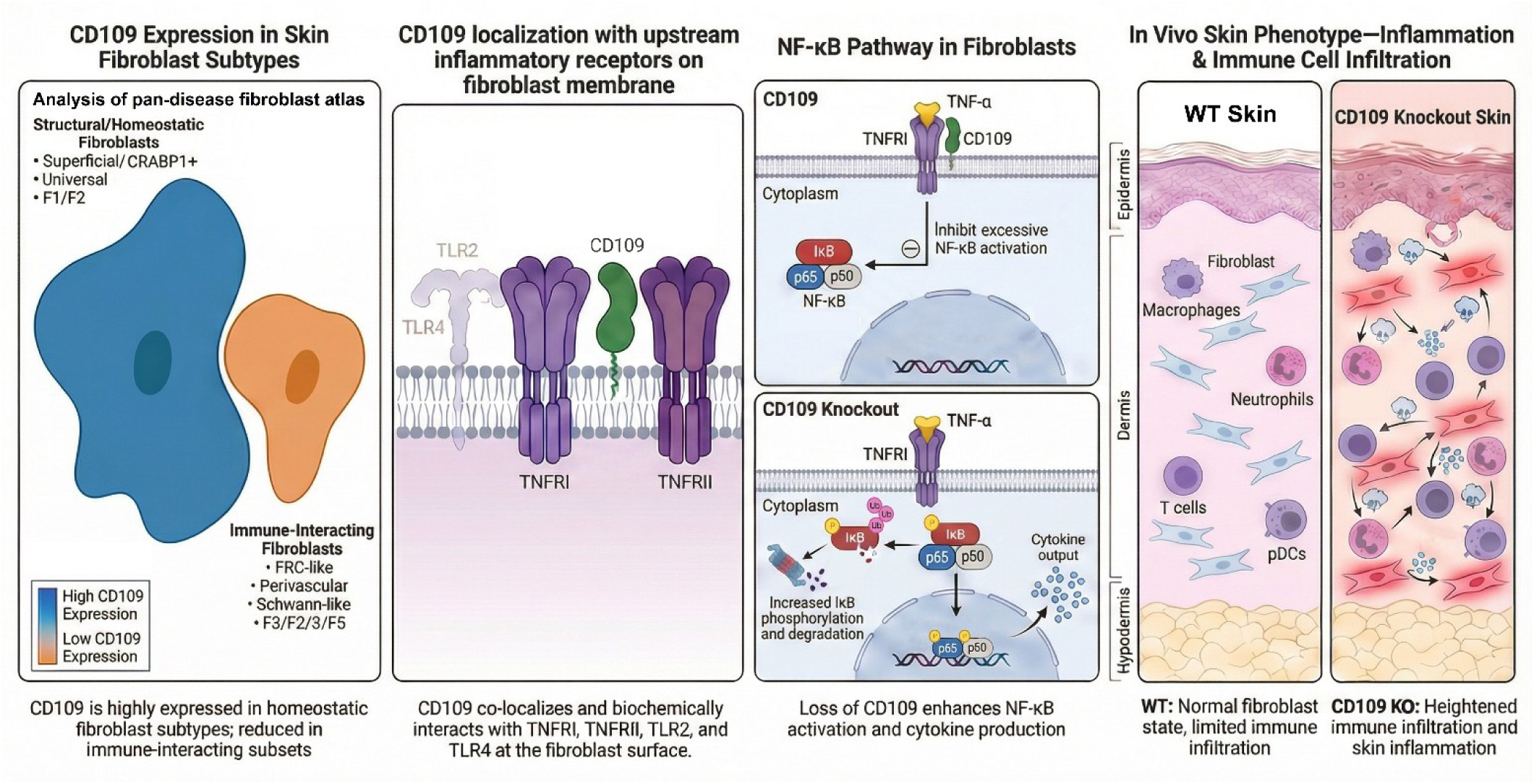

## Introduction

The skin is the largest organ of the human body and plays an active role in maintaining its overall physiological balance [1]. It is a multifunctional organ that connects the body to its environment, serving both as a protective barrier against pathogens, toxins, allergens, and chemicals and as a frontline immune organ populated by macrophages, neutrophils, dendritic cells, and T cells. [2]. Skin cells actively communicate with the immune system through the production of various cytokines and chemokines [2]. Dermal fibroblasts release cytokines, such as Interleukin 1 (IL-1), Tumor Necrosis Factor Alpha (TNF-α), IL-6, and Transforming Growth Factor Beta (TGF-β), which initiate and regulate inflammatory and repair responses following injury or infection [3, 4]. In addition to directing the recruitment and activation of immune cells at the site of skin stress and damage [5–7], fibroblasts work together with keratinocytes, immune cells, and endothelial cells to restore skin integrity through inflammation, cell proliferation, as well as extracellular matrix (ECM) deposition and remodeling [8, 9]. Dysregulation of these processes can lead to chronic wounds, fibrosis, or excessive scarring [10]. Prolonged inflammation with sustained TNF-α, IL-1β, and IL-6 expression can impair keratinocyte migration and reduce fibroblast proliferation, leading to a failure of re-epithelialization and persistent open wounds [11, 12]. Excessive TGF-β/Smad signaling maintains fibroblasts in an activated, myofibroblast state, causing excessive ECM deposition, increased dermal thickness, and progressive loss of tissue elasticity [13]. Together, these dysregulated inflammatory and fibrotic pathways underpin the pathogenesis of various skin diseases, including chronic non-healing ulcers, and pathological scarring as in hypertrophic scars, keloids, and systemic sclerosis (SSc/scleroderma) [14–17].

SSc is a rare autoimmune disease characterized by vasculopathy, chronic inflammation and progressive fibrosis of the skin and internal organs, including the heart, the lungs, the kidneys, and the gastrointestinal tract [18]. Its prevalence in North America varies, with reported ranges of approximately 13.5 to 44.3 cases per 100,000 people, and incidence estimates of 1.4 to 5.6 per 100,000 person-years. Currently, there is no cure for SSc, and treatment strategies largely rely on symptom management only. Immunosuppressive therapies like mycophenolate mofetil, methotrexate or cyclophosphamide are used to reduce disease activity and slow the progression of skin and lung fibrosis [19]. Antifibrotic agents, such as nintedanib, slow down the progression of SSc-associated interstitial lung disease (SSc-ILD) [20]. However, unlike SSc-ILD, no approved therapy currently exists to directly reverse or halt skin fibrosis in SSc. Therefore, a deeper understanding of the molecular mechanisms driving skin fibrosis is essential for the development of more effective therapeutic strategies that target the underlying causes of the disease.

The hallmark cellular signaling pathways involved in fibrotic conditions such as SSc are dominated by the fibrotic TGF-β and the inflammatory Nuclear Factor Kappa B (NF-κB) pathways [18, 21]. Dysregulated TGF-β signaling increases alpha smooth muscle actin (α-SMA) expression and stress fibre formation, shifting the dermal fibroblasts towards a myofibroblast phenotype [22]. These differentiated myofibroblasts significantly enhance ECM production and deposition mainly through canonical Smad2/3 signaling and via non-canonical pathways (MAPK, YAP/TAZ) [21–24]. Concurrently, the NF-κB pathway can influence the dermal fibroblasts by transitioning them toward an inflammatory phenotype in response to TNF-α and IL-1β [25]. Upstream of NF-κB activation, innate immune receptors such as Toll-like receptors (TLRs) and TNF receptors (TNFRs) play a critical role in sustaining inflammatory signaling in SSc. TLR4 is upregulated in lesional skin and lung tissue in SSc, where it co-localizes with myofibroblasts, macrophages, and vascular cells and correlates with disease progression [26]. Additionally, TLR2 expression is also increased in SSc, with rare functional variants associated with exaggerated IL-6 responses in patient-derived dendritic cells [27]. Via TNFR1 and TNFR2, TNF-α signaling can stimulate NF-κB, cytokine production and fibroblast survival, thus maintaining chronic inflammation in SSc [28]. These factors collectively create a pro-inflammatory environment that increases TGF-β-driven fibrosis [28, 29].

Although TGF-β and NF-κB signaling are known to be key drivers of fibrosis and inflammation in fibrotic disorders, these pathways are dynamically regulated at multiple levels, including by cell surface co-receptors. Our group has identified CD109 as a TGF-β co-receptor and negative regulator of fibrotic signaling in skin cells including scleroderma fibroblasts [30–35].

CD109 is a glycosylphosphatidylinositol (GPI)-anchored protein that binds to TGF-β1 and inhibits its downstream effects [32, 36]. CD109 also promotes the internalization and degradation of TGF-β receptors, reducing cellular sensitivity to TGF-β and limiting the intensity of downstream Smad-dependent signaling [37]. Our group has shown that epidermal overexpression of CD109 in mice attenuates TGF-β/Smad2/3 signaling, leading to reduced bleomycin-induced skin fibrosis as well as diminished inflammation and granulation tissue formation [38, 39]. Conversely, CD109 knockout (KO) mice display enhanced TGF-β/Smad2/3 signaling and skin fibrosis [40] accompanied by epidermal hyperplasia, impaired hair growth, and heightened immune cell infiltration [41]. While CD109 has been extensively studied in the context of TGF-β signaling, its role in modulating inflammatory signaling pathways, particularly the NF-κB pathway, remains uninvestigated. Our recent analysis of published studies on CD109 expression and function suggests that CD109 is likely a critical mediator of inflammatory responses across multiple tissues [42]. Additionally, we have shown that CD109 overexpression in the epidermis reduced immune cell infiltration (macrophages, neutrophils) in the skin [39, 43].

The present study establishes CD109 as a critical regulator of skin inflammation through modulation of the NF-κB pathway. Using both global and conditional CD109 knockout mouse models, we demonstrate that loss of CD109 markedly enhances NF-κB activation, increases pro-inflammatory cytokine expression, and promotes immune cell infiltration in the skin. Collectively, these findings identify CD109 as a key negative regulator of inflammatory signaling and underscore its pivotal role in maintaining cutaneous immune homeostasis.

## Results

### CD109 interacts with innate Toll-like and TNF receptors

To determine whether CD109 interacts with upstream innate and inflammatory receptors, we examined its spatial co-localization and biochemical interaction with TLR2, TLR4, TNFRI, and TNFRII. Immunofluorescence analysis revealed overlapping localization of CD109 with all four receptors (TLR2, TLR4, TNFRI, and TNFRII) in skin fibroblasts (**Figure 1A**). High-magnification images demonstrate their co-localization at the membrane and perimembranous regions of fibroblasts (**Figure 1A**). To validate these observations biochemically, we performed co-immunoprecipitation assays. Immunoprecipitation of CD109 pulled down TLR2 and TLR4, whereas immunoprecipitation of TNFRI and TNFRII pulled down CD109, as detected by immunoblotting (**Figure 1B-D**).

### CD109 deletion increases NF-κB activation and pro-inflammatory responses

To determine whether CD109 regulates the inflammatory NF-κB signaling pathway, we transfected human dermal fibroblasts with siRNA targeting CD109 to knock down (KD) its expression. After TNF-α treatment, Inhibitor of NF-κB α (IκBα) phosphorylation and degradation were significantly enhanced in CD109 KD fibroblasts (**Figure 2A-F**). CD109 KD also enhanced the activation of the NF-κB subunits, p50 and p65. Increased phosphorylation of the p50 (**Figure 2G-K**) and p65 (**Figure 2L-P**) subunits is observed in the CD109 KD fibroblasts after TNF-α treatment.

**Figure 2:**
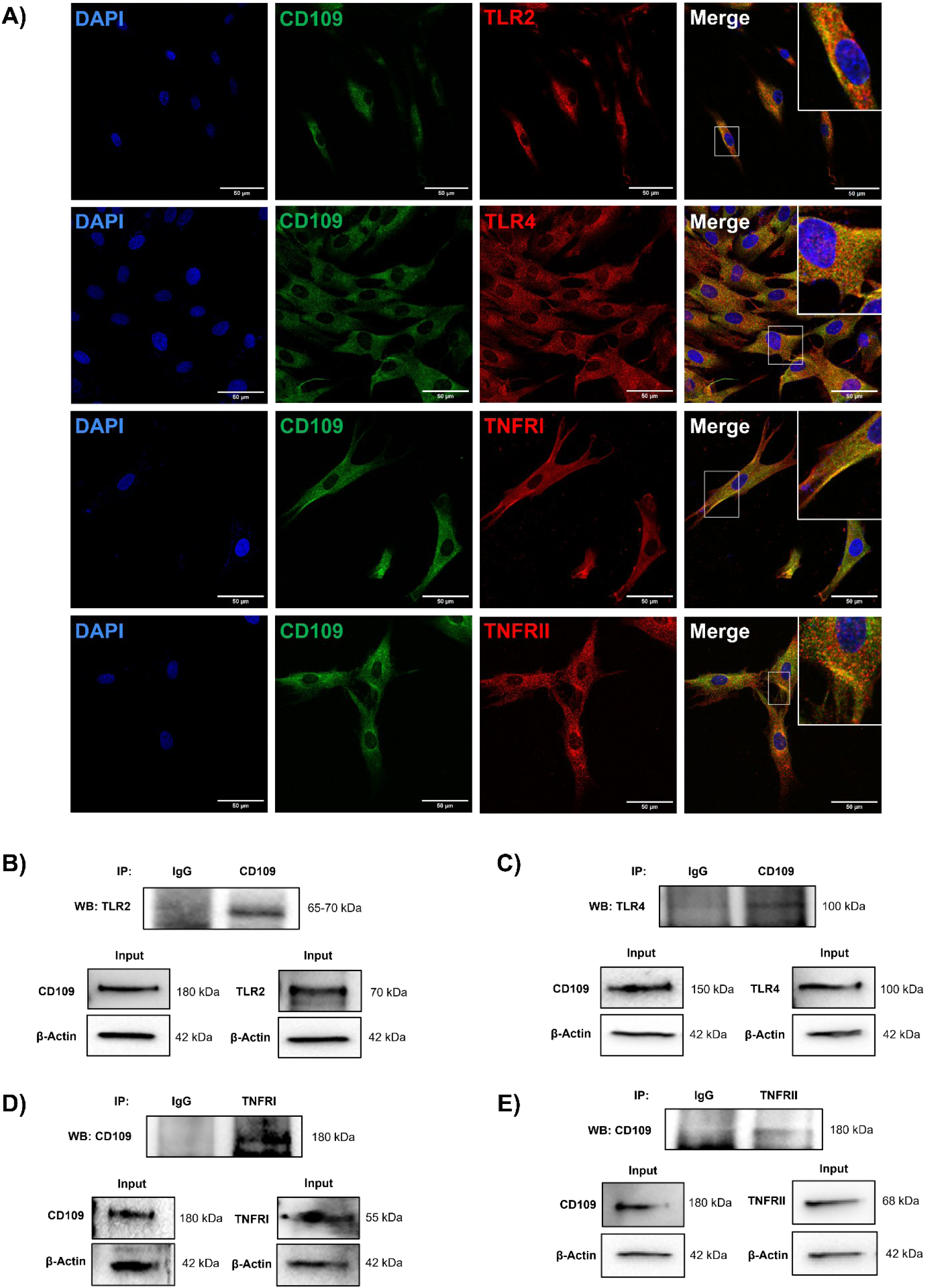
CD109 colocalizes and associates with TLR2/4 and TNFRI/II in skin fibroblasts. **A)** Representative immunofluorescence images of human dermal fibroblasts stained for CD109 (green), TLR2, TLR4, TNFRI, or TNFRII (red), with nuclei counterstained with DAPI (blue). Merged images show partial co-localization of CD109 with each receptor at the plasma membrane and perimembranous regions. Insets highlight areas of overlap between CD109 and the indicated receptors. Scale bars, 50 µm. **B)** Co-immunoprecipitation (Co-IP) assay through CD109 pulldown followed by immunoblotting for TLR2. **C)** Co-IP assay through CD109 pulldown followed by immunoblotting for TLR4. **D)** Co-IP assay through TNFRI pulldown followed by immunoblotting for CD109. **E)** Co-IP assay through TNFRII pulldown followed by immunoblotting for CD109.

To determine whether enhanced NF-κB activation following CD109 KD translates into altered inflammatory output, we assessed cytokine and chemokine expression in human dermal fibroblasts following CD109 depletion. Multiplex cytokine analysis revealed a broad alteration of inflammatory mediator expression in CD109 KD fibroblasts compared to controls (**Figure 3A**). CD109 KD fibroblasts exhibited relative increases in several pro-inflammatory cytokines, notably IL-6, IL-12p70, IL-15, and IL-16 (**Figure 3A**). In contrast, anti-inflammatory mediators were reduced, including IL-1 receptor antagonist (IL-1ra) and soluble TNFRII, while a relative increase in soluble TNFRI was observed (**Figure 3A**). To quantitatively validate these observations, cytokine concentrations were calculated from standard curves using linear regression and compared between conditions **(Supplementary Figure 1)**. This analysis demonstrated significant reductions in IL-1ra (p<0.001), soluble TNFRII (p<0.01), MIP-1δ (p<0.01), IL-7 (p<0.0001), MCP-1 (p<0.01), and TNF-α (p<0.001) concentrations in CD109 knockdown fibroblasts relative to controls, whereas IL-15 (p<0.05) concentrations were significantly increased following CD109 depletion (**Figure 3B-H**).

**Figure 3:**
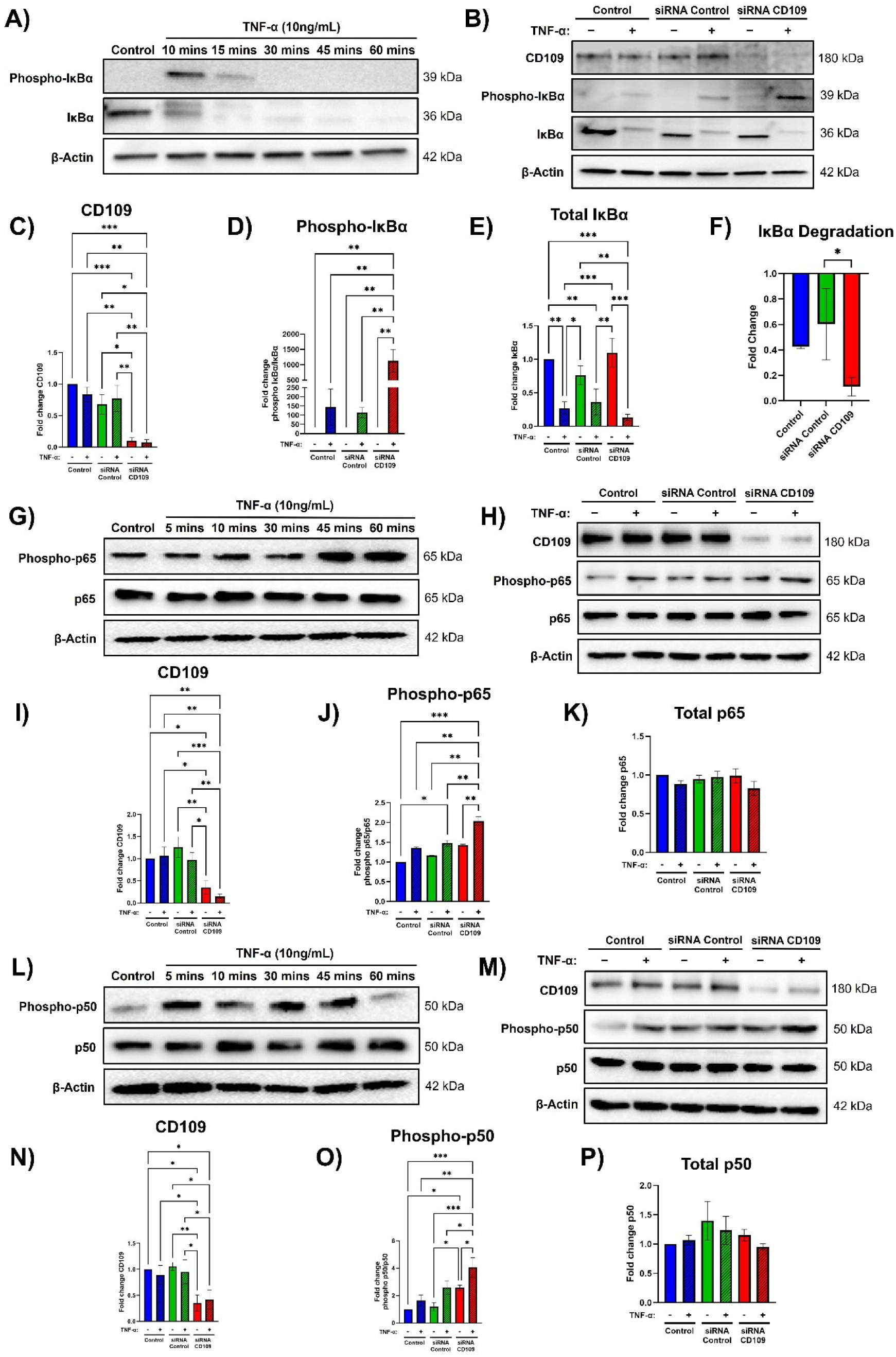
CD109 knockdown enhances TNF-α–induced NF-κB activation in human dermal fibroblasts. A–C) Immunoblot analysis of CD109, phosphorylated IκBα (phospho-IκBα), and total IκBα in human dermal fibroblasts transfected with control or CD109-targeting siRNA and treated with TNF-α, as indicated. CD109 knockdown efficiency is shown by reduced CD109 protein levels. **D)** Quantification of IκBα degradation following TNF-α stimulation, expressed as fold change relative to untreated control conditions. **E-G)** Immunoblot analysis and quantification of phosphorylated p65 (phospho-p65) and total p65 in control and CD109-knockdown fibroblasts following TNF-α treatment. **H-J)** Immunoblot analysis and quantification of phosphorylated p50 (phospho-p50) and total p50 in control and CD109-knockdown fibroblasts following TNF-α treatment. Protein band intensities were quantified by densitometry, normalized to loading controls, and expressed as fold change relative to untreated control samples. Data: Mean ± SEM (n=3 per blot). Statistical significance was determined using one-way ANOVA tests. (*p < 0.05; **p < 0.01; ***p < 0.001).

### Global and fibroblast-specific loss of CD109 leads to increased immune cell infiltration and inflammation in mouse skin

To assess the effect of CD109 deletion on skin inflammation, we used two complementary *in vivo* mouse models: a global CD109 KO model and a tamoxifen-inducible fibroblast-specific CD109 KO model **(Supplementary Figure 2A-B)**. The use of these two models provides an important advantage. The global knockout of CD109 defines its overall physiological and pathological functions at the whole-organism level, and because CD109 is deleted in all tissues, this model reveals systemic effects and interactions between different cell types. However, developmental compensation and contributions from multiple cell populations can obscure the specific cellular mechanisms underlying the observed phenotype. In the fibroblast-specific inducible knockout model, as CD109 is deleted in fibroblasts while preserving expression in other cell types, developmental confounders are minimized, allowing direct assessment of fibroblast-intrinsic functions, even though the model does not capture contributions from other cell types. The fibroblast-specific KO model allowed determining whether CD109 deletion specifically in fibroblasts is sufficient to promote an inflammatory phenotype. This approach is particularly relevant because fibroblasts are increasingly recognized not only as structural cells, but also as active regulators of immune responses in the skin microenvironment. Thus the combined use of global and fibroblast-specific inducible knockout models provides complementary insights, enabling discrimination between the overall physiological role of CD109 and its cell-autonomous functions in fibroblasts.

Immunohistochemical analysis for immune cell markers was performed on the skin of both models and their wild-type (WT) counterparts. Global CD109 KO mouse skin exhibited significantly reduced levels of IκBα (p<0.0001) compared to WT skin (**Figure 4A**). Moreover, the global CD109 KO skin showed significant increases in immune cell markers, including macrophages (F4/80, p<0.0001), neutrophils (Ly-6G, p<0.0001), T cells (CD3, p<0.0001), and plasmacytoid dendritic cells (pDCs; BST2, p<0.0001), compared to WT skin (**Figure 4B-E**). Similarly, fibroblast-specific CD109 KO mouse skin revealed a significant increase in those same immune cell markers (**Figure 5**).

**Figure 4:**
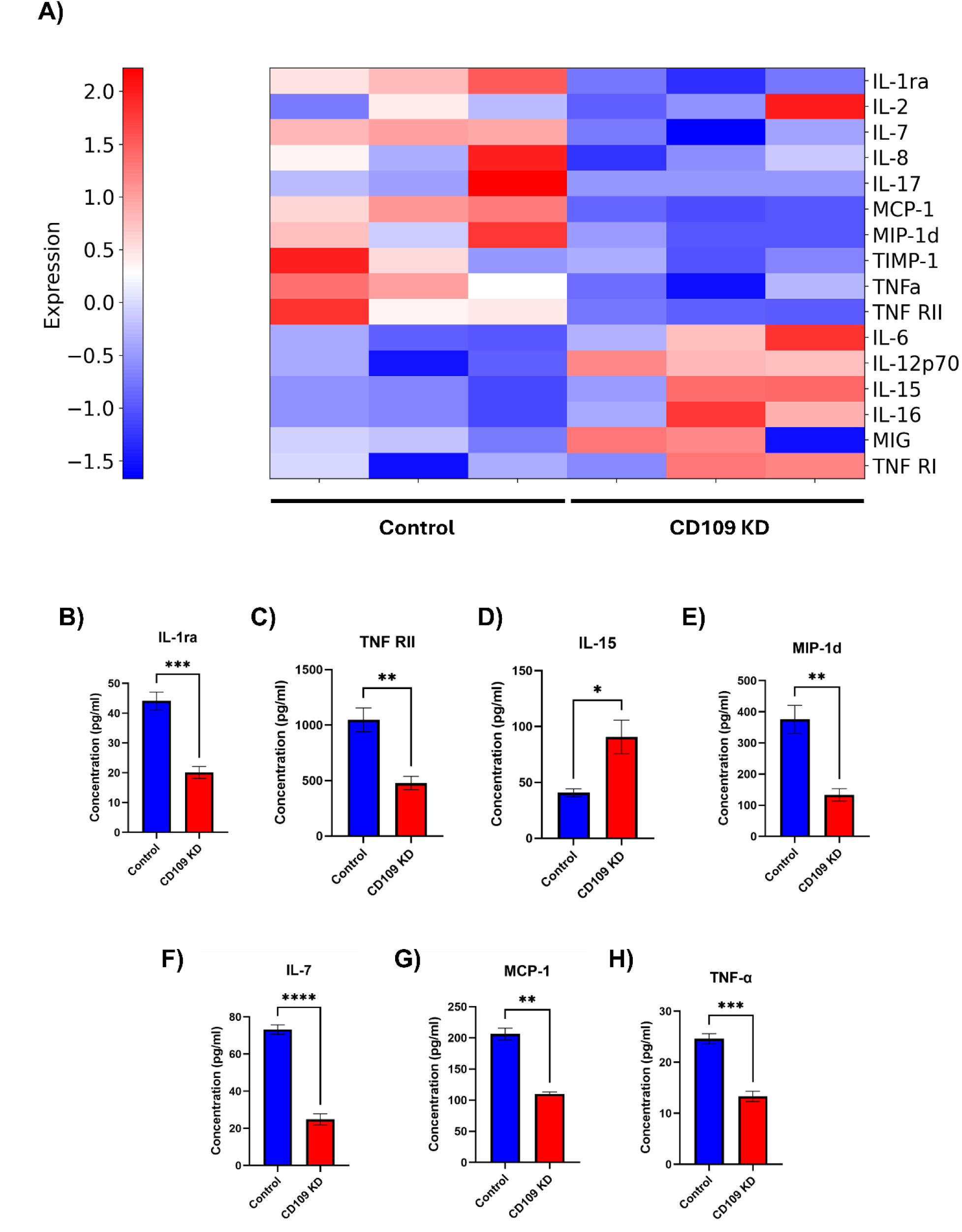
CD109 deletion results in a selective remodelling of the fibroblast inflammatory secretome. **A)** Heatmap showing relative expression levels of inflammatory cytokines, chemokines, and TNF receptor components in conditioned media from control and CD109-knockdown human dermal fibroblasts. Values are scaled per analyte to highlight relative differences across conditions. Red indicates higher expression, and blue indicates lower expression. **B-H)** Quantification of significant cytokines and chemokines in conditioned media from control and CD109-knockdown (KD) fibroblasts measured by multiplex immunoassay. Shown are IL-1 receptor antagonist (IL-1ra), soluble TNF receptor II (TNF RII), IL-15, MIP-1d, IL-7, MCP-1, and TNF-α. Data are expressed as concentration (pg/mL). Data: Mean ± SEM (n=3). Statistical significance was assessed using Welch’s t test (*p < 0.05; **p < 0.01; ***p < 0.001; ****p < 0.0001).

**Figure 5:**
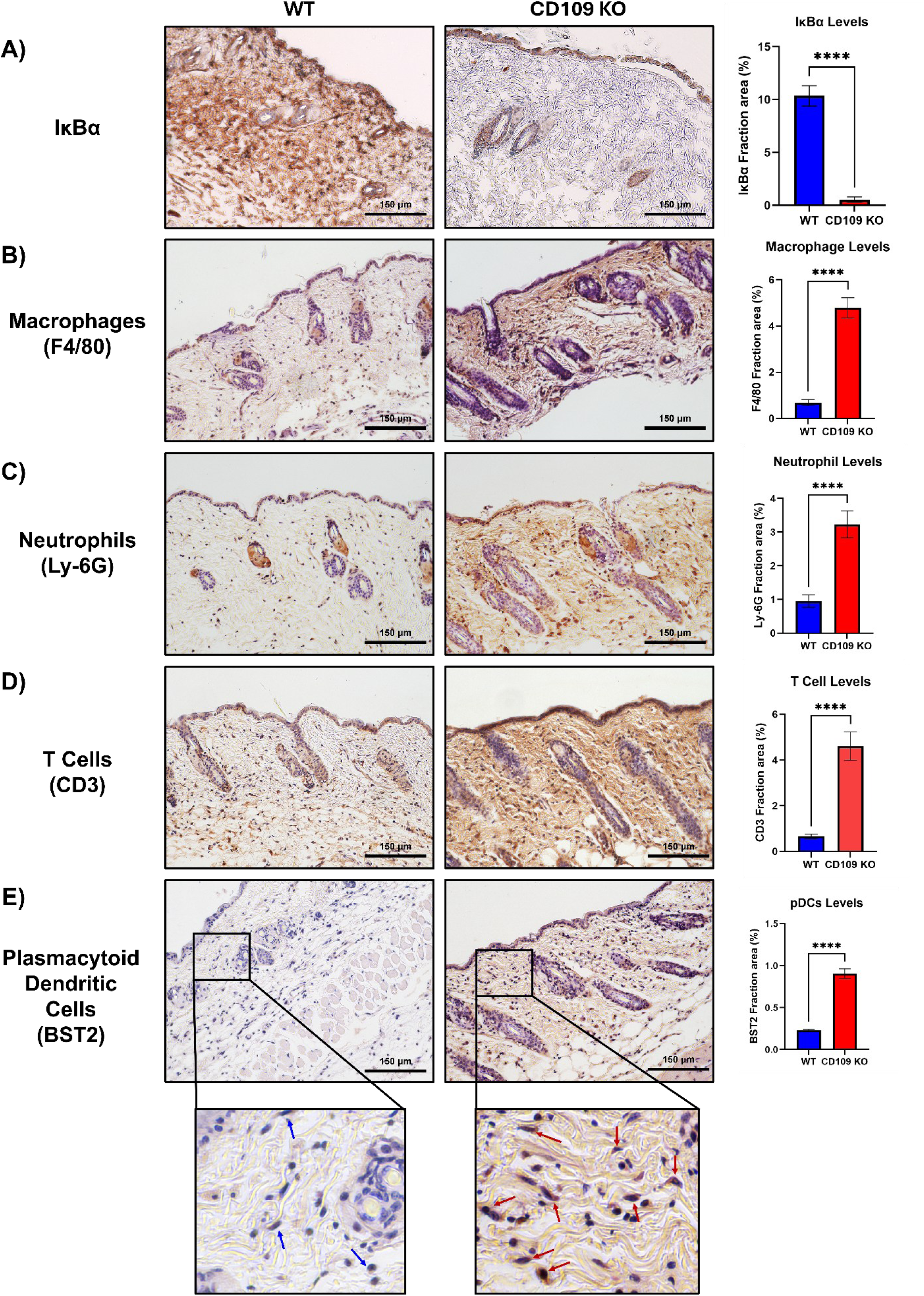
Global CD109 KO promotes immune cell infiltration in mouse skin. **A)** Representative immunohistochemical staining of IκBα in dorsal skin sections from WT and global CD109 KO mice. Quantification of the IκBα-positive area (% of tissue area) is shown to the right. **B)** Immunohistochemical staining for macrophages (F4/80) in WT and CD109 KO skin. Quantification of the F4/80-positive area is shown to the right. **C)** Immunohistochemical staining for neutrophils (Ly6G) in WT and CD109 KO skin. Quantification of the Ly6G-positive area is shown to the right. **D)** Immunohistochemical staining for T cells (CD3) in WT and CD109 KO skin. Quantification of the CD3-positive area is shown to the right. **E)** Immunohistochemical staining for plasmacytoid dendritic cells (pDCs; BST2) in WT and CD109 KO skin. Quantification of the BST2-positive area is shown to the right. Higher-magnification images of dermal regions from WT and CD109 KO skin highlighting individual pDCs (arrows). Data: Mean ± SEM (n=6). Statistical significance was assessed using one-way ANOVA (****p < 0.0001). Scale bars, 150 μm.

**Figure 5:**
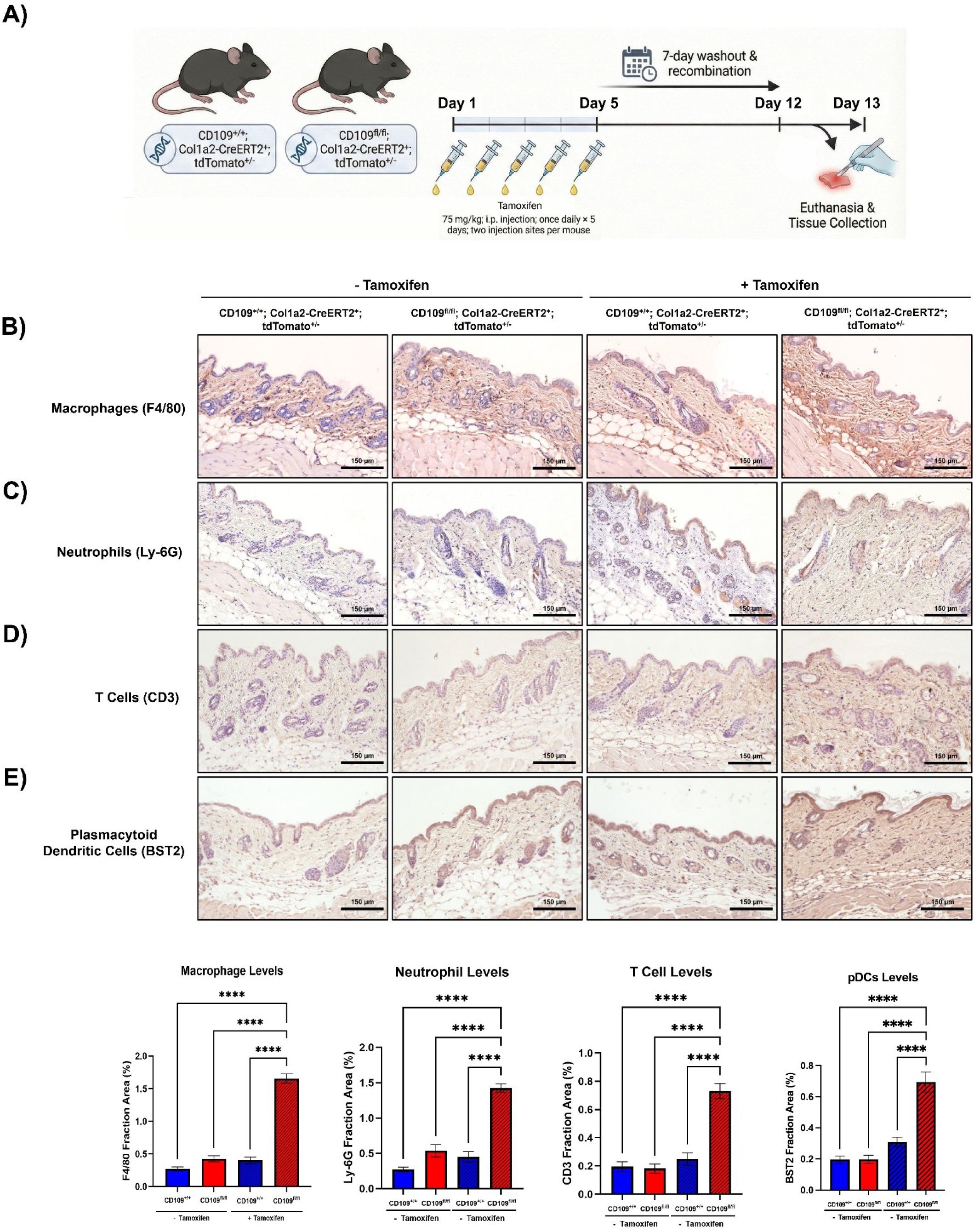
Conditional fibroblast-specific CD109 KO promotes immune cell infiltration in mouse skin. **A)** Experimental workflow. Image generated using *FigureLabs.*ai. **B)** Immunohistochemical staining for macrophages (F4/80). Quantification of the F4/80-positive area is shown to the right. **C)** Immunohistochemical staining for neutrophils (Ly6G). Quantification of the Ly6G-positive area is shown to the right. **D)** Immunohistochemical staining for T cells (CD3). Quantification of the CD3-positive area is shown to the right. **E)** Immunohistochemical staining for plasmacytoid dendritic cells (pDCs; BST2). Quantification of the BST2-positive area is shown to the right. Data: Mean ± SEM (n=3 per group). Statistical significance was assessed using one-way ANOVA (****p < 0.0001). Scale bars, 150 μm.

### CD109 expression is inversely associated with immune-associated fibroblast subtypes and inflammatory pathway activity in SSc

Analysis of data (curated by publicly available Human Protein Atlas) representing 34 single cell and single nuclear RNA sequencing data sets show that across the human body, CD109 displays cell-type-restricted expression with enrichment in endocrine, immune, and mesenchymal cells (**Figure 6A**). Among mesenchymal compartments, skin fibroblasts exhibit the highest levels of CD109 expression (**Figure 6A**). The projection of CD109 expression onto the pan-disease fibroblast UMAP reveals marked differences between fibroblast subtypes rather than a uniform expression across all fibroblasts (**Figure 6B, 6C)**. CD109 expression is most prominent in Superficial (F1), Superficial CRABP1+ (F1) fibroblasts, and Fascia-like Myofibroblasts (F8), while lower expression is observed in Perivascular (F2/3), FRC-like (F3), and NGFR+ (F5) fibroblasts (**Figure 6C, 6E).** SSc fibroblasts mapped onto the pan-disease fibroblast atlas are predominantly distributed in Universal (F2, 25.9%), and Superficial (F1, 14.7%) fibroblast subtypes, with additional representation in FRC-like (F3, 13.6%), DS_DPEP1+ (F4, 11.2%), and perivascular (F2/3, 9.9%) populations (**Figure 6D, 6F)**. In contrast, SSc fibroblasts are present at low frequency in the inflammatory myofibroblasts (F6, 0.4%) and other immune-associated fibroblast subsets, including Schwann-like fibroblasts (F5: RAMP1+, 1.9%; F5: NGFR+, 3.0%) (**Figure 6D, 6F)**. Despite heterogeneity in subtype distribution, CD109 expression is broadly maintained across SSc fibroblast subtypes. Quantification of CD109 expression across Healthy and SSc fibroblast subtypes shows the higher mean expression across all SSc clusters compared to healthy clusters (**Figure 6G).** The highest fraction of CD109-positive cells in Universal (F2, 59.2%), Superficial (F1, 58.2%), Superficial CRABP1+ (F1, 53.4%), Fascia-like myofibroblast (F8, 52.9%), and Myofibroblast (F7, 48.1%) populations (**Figure 6G**). CD109 expression is reduced in SSc fibroblasts present in clusters associated with immune signaling, including FRC-like fibroblasts (F3), perivascular fibroblasts (F2/3), and Schwann-like fibroblasts (F5: RAMP1+, F5: NGFR+) (**Figure 6G**). Consistent with this distribution, Gene Set Variation Analysis (GSVA) performed on fibroblast pseudobulk samples highlights a significant downregulation of inflammatory and immune pathways in SSc fibroblasts, including the hallmark inflammatory response pathway and the TNF-α signaling pathway via NF-κB (**Figure 6H-I**).

**Figure 6:**
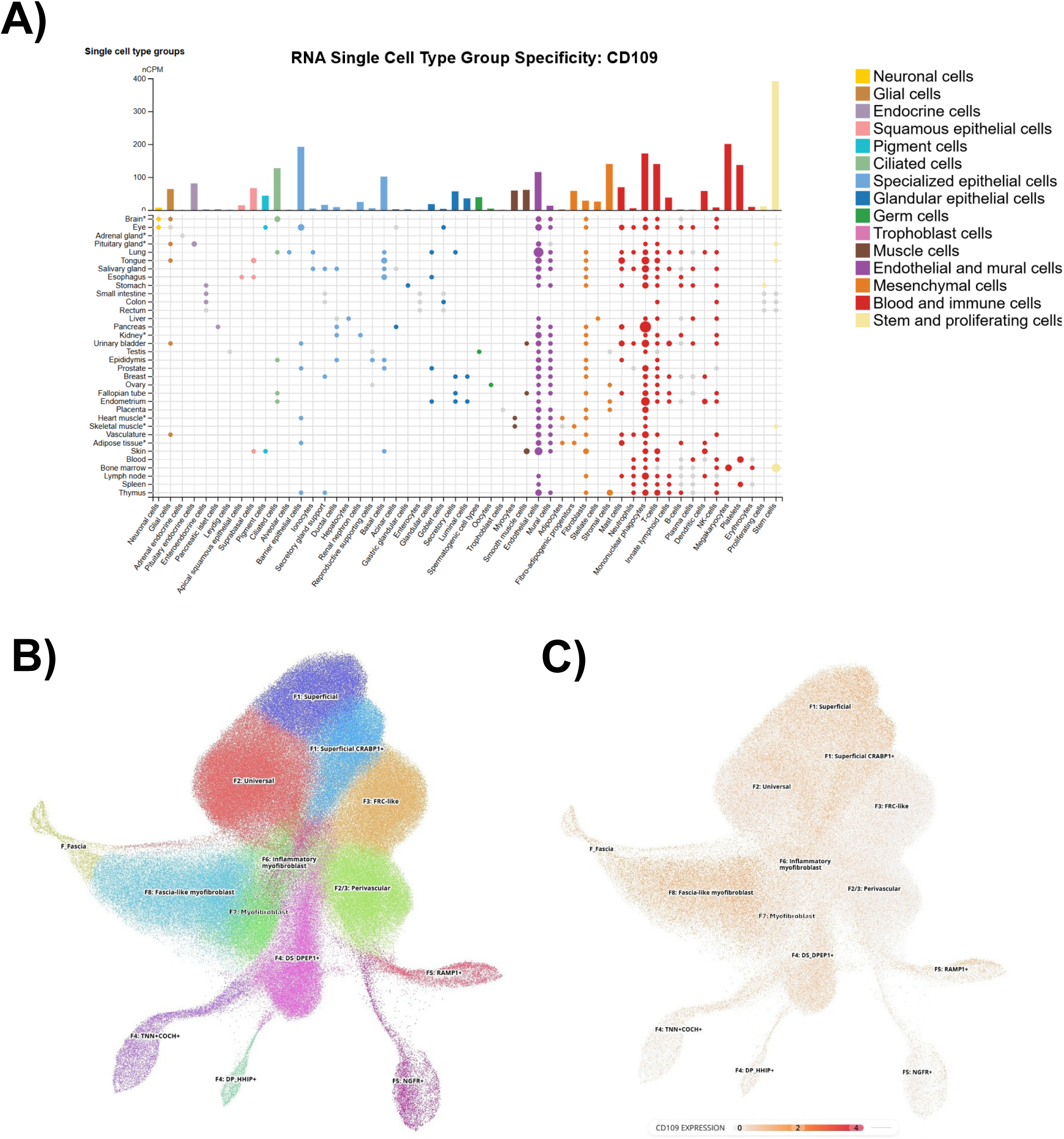
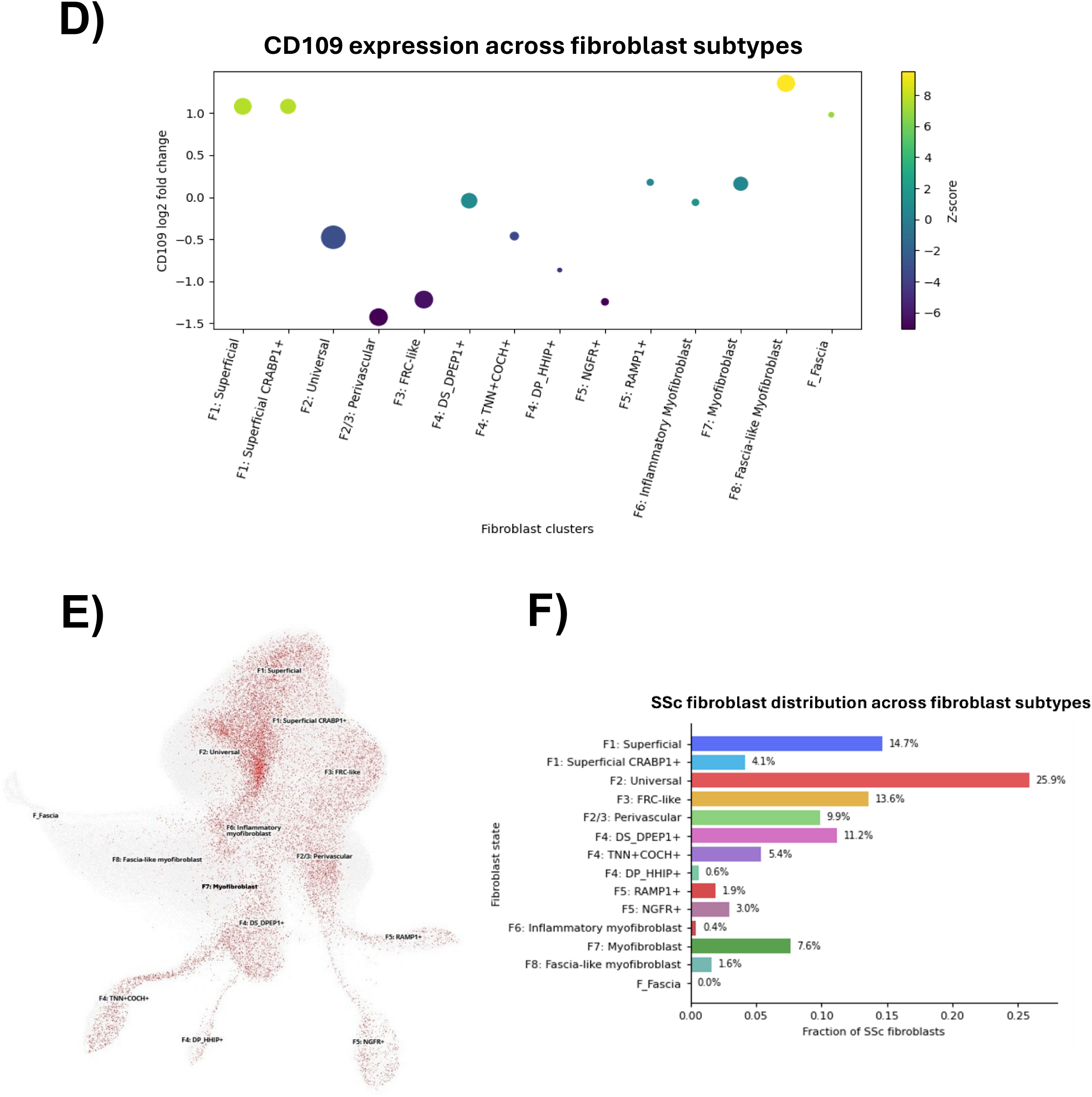
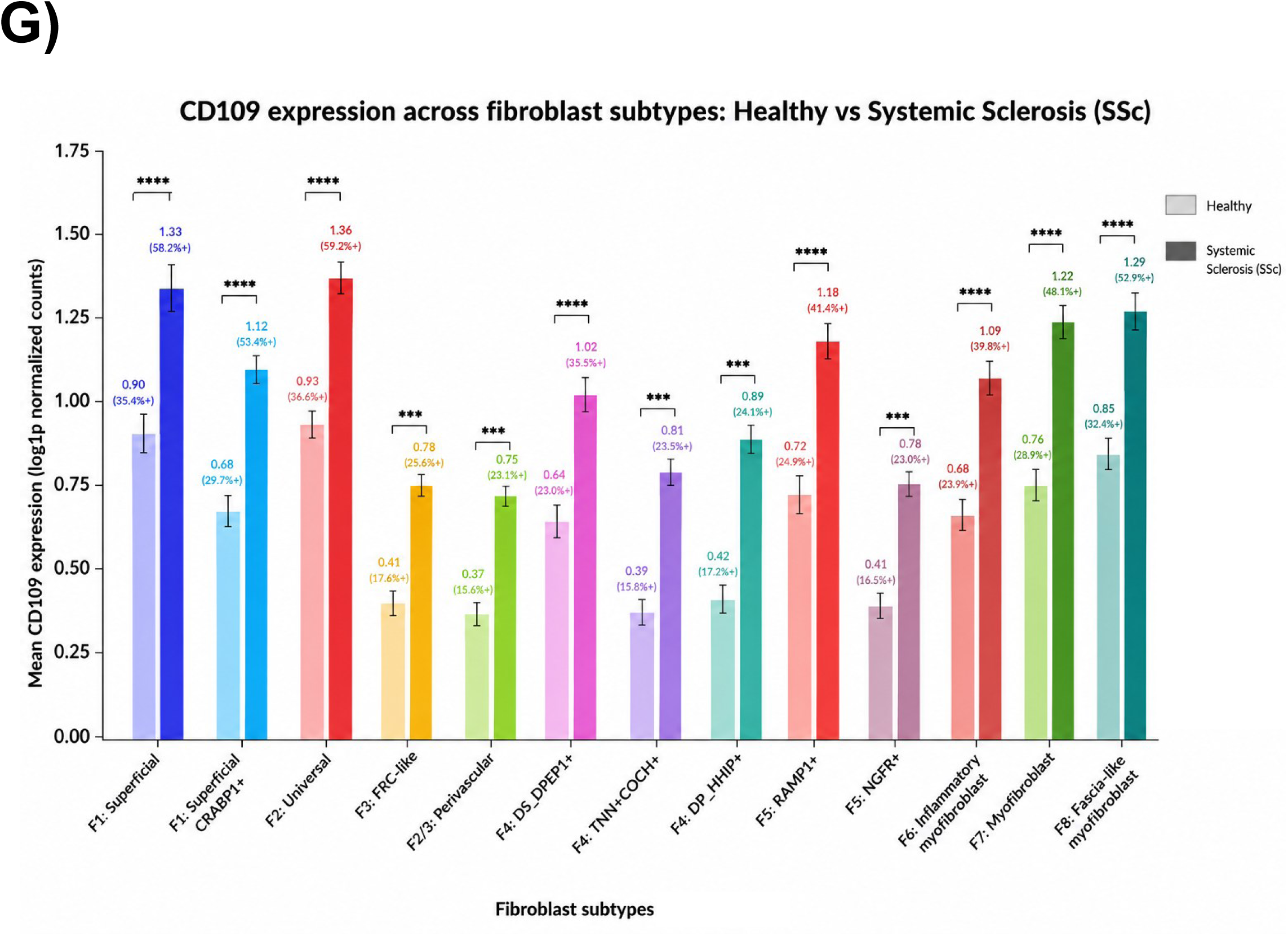
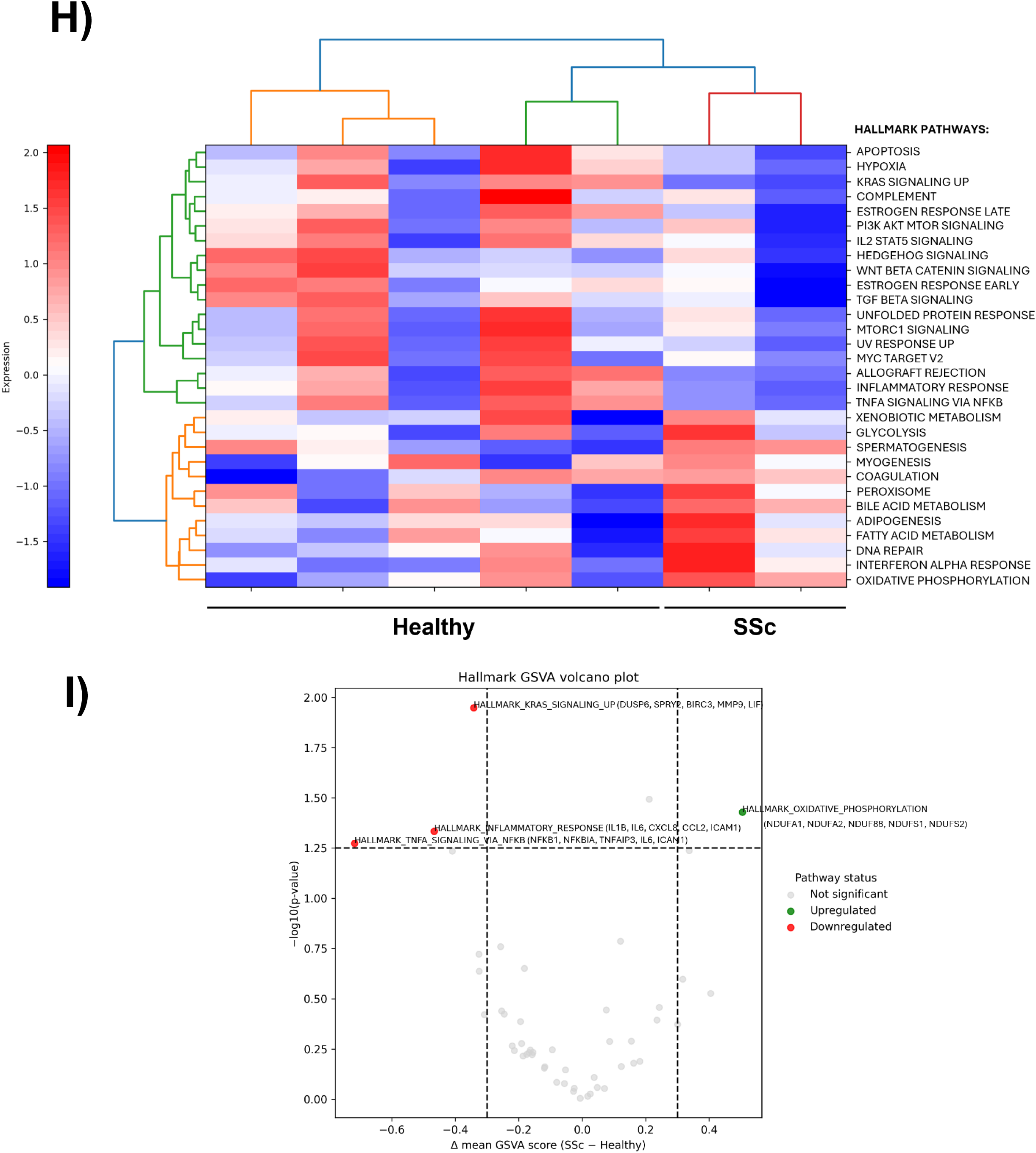
CD109 expression is enriched in structural fibroblast subtypes and reduced in immune-interacting fibroblasts in SSc. **A)** Cell-type-specific CD109 expression across the human body, derived from 34 publicly available single-cell and single-nucleus RNA sequencing datasets curated by the Human Protein Atlas. **B**) Uniform Manifold Approximation and Projection (UMAP) of the pan-disease skin fibroblast atlas, showing fibroblast subtypes identified across skin fibrosis, inflammation, and cancer. **C)** Projection of CD109 expression onto the pan-disease fibroblast UMAP, demonstrating heterogeneous expression across fibroblast subtypes. **D)** Bubble plot showing CD109 expression across pan-disease fibroblast subtypes, where dot size represents the fraction of CD109-positive cells and color indicates mean expression levels. **E)** UMAP highlighting SSc fibroblasts (n = 27,444 cells) mapped onto the pan-disease fibroblast atlas. **F)** Distribution of **SSc fibroblasts** across fibroblast subtypes defined in the pan-disease atlas. **G)** Mean CD109 expression across Healthy and SSc fibroblast subtypes. Bars represent mean log1p-normalized CD109 expression within each subtype among Healthy and SSc fibroblasts. Percentages indicate the fraction of CD109-positive cells (expression > 0) within each subtype. **H)** Heatmap of Hallmark Gene Set Variation Analysis (GSVA) scores across fibroblast pseudobulk samples derived from healthy vs SSc skin using the pan-disease fibroblast atlas. Color intensity reflects relative pathway activity. **I)** Volcano plot (data shown in H) demonstrating differential Hallmark pathway activity between SSc vs healthy fibroblast pseudobulk samples. The x-axis represents the difference in mean GSVA scores (SSc minus healthy), and the y-axis shows statistical significance (−log10 p-value). Vertical dashed lines indicate thresholds for biologically meaningful enrichment (|Δ GSVA| ≥ 0.3), and the horizontal dashed line indicates statistical significance (p < 0.05).

## Discussion

CD109 is known as a major regulator of skin fibrosis in SSc [30]. However, skin inflammation is also a major contributor to morbidity in SSc and is largely driven by NF-κB-dependent signaling[18]. Our results extend the role of CD109 beyond fibrosis and identify it as a negative regulator of fibroblast inflammatory signaling. CD109 seems to act as a receptor-associated regulator of inflammatory signaling. Immunofluorescence and co-immunoprecipitation analyses demonstrate CD109 interaction with TLR2, TLR4, TNFRI, and TNFRII. The partial co-localization observed by immunofluorescence (i.e., CD109 overlaps with TLR2/4 and TNFRI/II in discrete regions, not uniformly across the membrane) is consistent with the idea that these proteins are organized into specialized membrane microdomains (nanodomains) rather than diffusely distributed. GPI-anchored proteins are well known to concentrate signaling receptors and adaptor proteins, thereby shaping receptor responsiveness [44, 45], raising the possibility that CD109 acts as a spatial organizer that helps assemble or stabilize receptor signaling hubs. CD109 could influence inflammatory signaling by modulating receptor clustering, residence time at the plasma membrane, or endocytic trafficking. In fact, intracellular staining of CD109 and TLR/TNFR receptors, though modest, was observed, suggesting that CD109 may also be involved in receptor trafficking rather than acting exclusively at the plasma membrane. These findings could support a model in which CD109 constrains receptor organization or trafficking, such that its loss enhances effective receptor signaling and amplifies NF-κB–dependent inflammatory responses. Our finding that CD109 KD enhanced IκBα phosphorylation and degradation as well as increased p50/p65 phosphorylation, highlight overactivation of the NF-κB pathway. CD109 KD caused a marked shift in fibroblast cytokine profiles. The heatmap analysis revealed relative increases in several NF-κB–linked pro-inflammatory cytokines, including IL-6, alongside reductions in anti-inflammatory mediators such as IL-1ra and soluble TNFRII. Consistent with the complexity of NF-κB-driven inflammatory processes, CD109 KD did not result in a uniform upregulation of all pro-inflammatory cytokines. We observed reduced levels of TNF-α, MCP-1, and MIP-1δ. One potential explanation of this suppression is the known role of CD109 as a negative regulator of TGF-β signaling. Deletion of CD109 increased TGF-β activity, which can exert anti-inflammatory effects [46, 47]. Furthermore, the increase in soluble TNFRI, a marker of heightened inflammatory signaling in SSc [48, 49], can explain the decrease in TNF-α levels. Soluble TNFRI can act as a decoy receptor and limit TNF-α bioavailability [50]. These findings indicate that CD109 deletion can lead to a reprogramming of inflammatory signaling, shaped by the intersection of the NF-κB, TGF-β, and TNF receptor-mediated mechanisms.

The *in vivo* data further support the role of CD109 in regulating skin inflammation. Both the global and fibroblast-specific CD109 knockout mice exhibit a marked increase in immune cell infiltration (macrophages, neutrophils, T cells, and pDCs) in the skin. A major strength of this study is the use of the two complementary *in vivo* models. The global CD109 knockout model shows the overall consequence of complete CD109 loss in the skin, while the tamoxifen-inducible fibroblast-specific knockout model provides a more mechanistic approach by determining whether CD109 loss specifically in fibroblasts is sufficient to drive inflammation. The infiltration of plasmacytoid dendritic cells (pDCs) in the mouse skin mirrors the pathogenesis of SSc, where they also infiltrate the skin of SSc patients [51]. The similar phenotype observed in both mouse models indicates that fibroblast-intrinsic CD109 helps suppress inflammatory signaling and immune cell recruitment. Importantly, the fibroblast-specific model strengthens the novelty of our approach by showing that fibroblast-derived CD109 is sufficient to regulate the inflammatory skin microenvironment in vivo, rather than this effect being only a secondary consequence of global CD109 deletion. Without CD109, fibroblasts seem to create a permissive inflammatory niche.

Single-cell analyses across the pan-disease fibroblast atlas (https://celltype.info/project/613) [52] demonstrate that CD109 expression is preferentially enriched in SSc fibroblasts compared to healthy fibroblasts, more specifically in structural and homeostatic SSc fibroblast subtypes, including Universal (F2) and Superficial (F1) fibroblasts, which are not primarily specialized for immune interaction. In contrast, SSc fibroblast subsets associated with immune cell interactions, such as the FRC-like fibroblasts (F3), perivascular (F2/3), and Schwann-like fibroblasts (F5), consistently exhibit lower CD109 expression. It seems like CD109 is selectively reduced in the above-mentioned clusters characterized by immune engagement, indicating that its expression aligns with fibroblast cell states in SSc. This observation raises the possibility that CD109 may buffer fibroblasts against inflammatory polarization and is consistent with immunohistochemical analyses showing higher immune cell infiltration in CD109 KO, implying that loss of CD109 may promote fibroblast states that facilitate immune engagement. Additionally, GSVA performed on fibroblast pseudobulk samples shows reduced activity of canonical inflammatory pathways, including TNF-α signaling via NF-κB, in all SSc fibroblasts, which have high CD109 expression compared to healthy fibroblasts.

Importantly, several limitations should be considered when interpreting our findings. First, the human single-cell analyses are correlative and do not establish a causal relationship between CD109 expression and inflammatory pathway activity. In addition, the pseudobulk GSVA comparing SSc and healthy fibroblasts was not stratified by CD109 expression because of the limited number of SSc patient samples (n=2); therefore, these findings remain inconclusive. Furthermore, although immunofluorescence and co-immunoprecipitation assays demonstrate spatial proximity and biochemical association between CD109 and upstream receptors of NF-κB signaling, these approaches do not distinguish direct receptor binding. Finally, while mouse models provide robust in vivo evidence for a fibroblast-intrinsic role of CD109 in modulating skin inflammation, species-specific differences may limit direct translation to human disease.

To further strengthen these findings and better define the relationship between CD109 expression and fibroblast inflammatory states in SSc, future studies using CD109-stratified single-cell analyses will be important. Moreover, high-resolution approaches to investigate protein–protein interactions, such as cryo-electron microscopy, could help determine whether CD109 directly interacts with TLR2/4 and TNFRI/II. In parallel, live-cell imaging and trafficking-focused approaches will enable the determination of whether CD109 regulates the internalization, recycling, or endosomal routing of TLR2/4 and TNFRI/II. Collectively, our findings identify CD109 as an intrinsic anti-inflammatory regulator of fibroblasts that limits NF-κB-dependent signaling and maintains fibroblast states with limited immune engagement in the skin.

## Materials and Methods

### Single-cell RNA sequencing analysis of datasets for CD109 cell-type specificity across human tissues

To assess the cell-type specificity of CD109 expression across the human body, we integrated curated single-cell transcriptomic datasets with cell-type group specificity annotations from the Human Protein Atlas (HPA). The HPA single-cell resource aggregates uniformly processed single-cell RNA sequencing datasets and provides standardized annotations of gene expression enrichment across major human cell types and tissues (https://www.proteinatlas.org). In total, 34 independent single-cell RNA sequencing datasets were analyzed, retrieved from the Single Cell Expression Atlas, Human Cell Atlas, Gene Expression Omnibus (GEO), EMBL-EBI BioStudies, and Tabula Sapiens, as incorporated within the HPA single-cell framework. Normalized expression values were aggregated at the cell-type group level, and CD109 expression specificity across major cell-type categories was visualized using the HPA cell-type group specificity framework (**Figure 6A**). To analyze CD109 expression across skin fibroblast subtypes, we used the pan-disease fibroblast atlas of skin fibrosis, inflammation, and cancer, which provides a unified fibroblast annotation framework across multiple disease contexts [52]. This atlas was originally described by Steele et al. [52] and integrates fibroblasts from inflammatory, fibrotic, and malignant skin conditions into a common reference space (https://celltype.info/project/613). Cells annotated as SSc in the Patient_status metadata were extracted for downstream analyses, yielding 27,444 SSc fibroblasts. SSc fibroblasts were projected onto the reference UMAP embedding to assess their distribution across fibroblast subtypes (**Figure 6D**). Pathway activity was quantified using Gene Set Variation Analysis (GSVA) [53] with Hallmark gene sets from the Molecular Signatures Database (MSigDB) (https://www.gsea-msigdb.org/gsea/msigdb). Volcano plots and bar graphs were generated using custom Python scripts using Matplotlib and Seaborn, while single-cell analyses were performed with Scanpy [54] and pathway-level analyses were conducted using GSVA-based workflows.

### Generation of Mouse Colonies and Tamoxifen Induction

Two mouse colonies were used in this study: a global CD109 knockout (KO) colony and a tamoxifen-inducible, fibroblast-specific CD109 conditional KO colony. The global CD109 KO mouse colony (n=6, 3 males and 3 females) was obtained from Dr. Masahide (Department of Pathology, Nagoya University Graduate School of Medicine, Nagoya, Japan) [55]. Mice were maintained on a C57BL/6 background. For experiments involving the global KO model, wild-type (WT) littermates derived from heterozygous breeding were used as controls (n=6, 3 males and 3 females). The conditional CD109 KO colony (CD109^fl/fl^; Col1a2-CreERT2^+^; tdTomato^+/-^; n=6, 3 per group, 2 males and 4 females) was generated by crossing CD109^fl/fl^ mice (generated at the McGill Integrated Core for Animal Modeling (MICAM), Rosalind and Morris Goodman Cancer Institute at McGill University) with Col1a2-CreERT2^+^; tdTomato^+/+^ mice. Control animals for Cre-lox experiments consisted of CD109^+/+^; Col1a2-CreERT2^+^; tdTomato^+/-^ (n=6, 3 per group, 2 males and 4 females) mice. Both conditional knockout and control mice were treated with tamoxifen or vehicle, as indicated, to control for potential effects of Cre expression and tamoxifen administration. Tamoxifen (Sigma-Aldrich) was dissolved in corn oil at a concentration of 20 mg/mL and administered to mice aged 6–8 weeks by intraperitoneal injection at a dose of 75 mg/kg body weight once daily for five consecutive days. Two injection sites were used per mouse. Following the final injection, mice were allowed a 7-day washout and recombination period prior to tissue collection. Mice were euthanized on day 13, and skin samples were harvested for histological and biochemical analyses. Throughout treatment and post-injection periods, animals were monitored for adverse effects.

### Genotyping of CD109 Mice

Genomic DNA was extracted from mouse ear punch biopsies by alkaline lysis. Tissue samples were incubated in 50 mM NaOH at 96°C for 1h. vortexed intermittently and neutralized with 1 M Tris-HCl (pH 8.0). DNA extracts were stored at −20°C until use. Genotyping was performed by PCR amplification using allele-specific primer sets (**Table 1**) to detect the Col1a2-CreERT2 transgene, tdTomato reporter, 5′ and 3′ LoxP sites flanking the *Cd109* locus, and the global CD109 knockout allele **(Supplementary Figure 3)**. PCR reactions were carried out using the OneTaq Quick-Load 2× Master Mix, according to the primer set used. PCR products were resolved on 1% or 3% agarose gels containing SYBR Safe and visualized under UV illumination. The presence of the Col1a2-CreERT2 transgene was confirmed by detection of a ∼1300 bp amplicon. The tdTomato reporter allele was identified by PCR fragments of 196 bp (tdTomato) and 297 bp (WT). Correct insertion of 5′ and 3′ LoxP sites was verified by amplification of fragments of approximately 300 bp. For the global CD109 knockout model, PCR amplification yielded bands of 205 bp for the wild-type allele and 603 bp for the mutant allele.

**Table 1.**
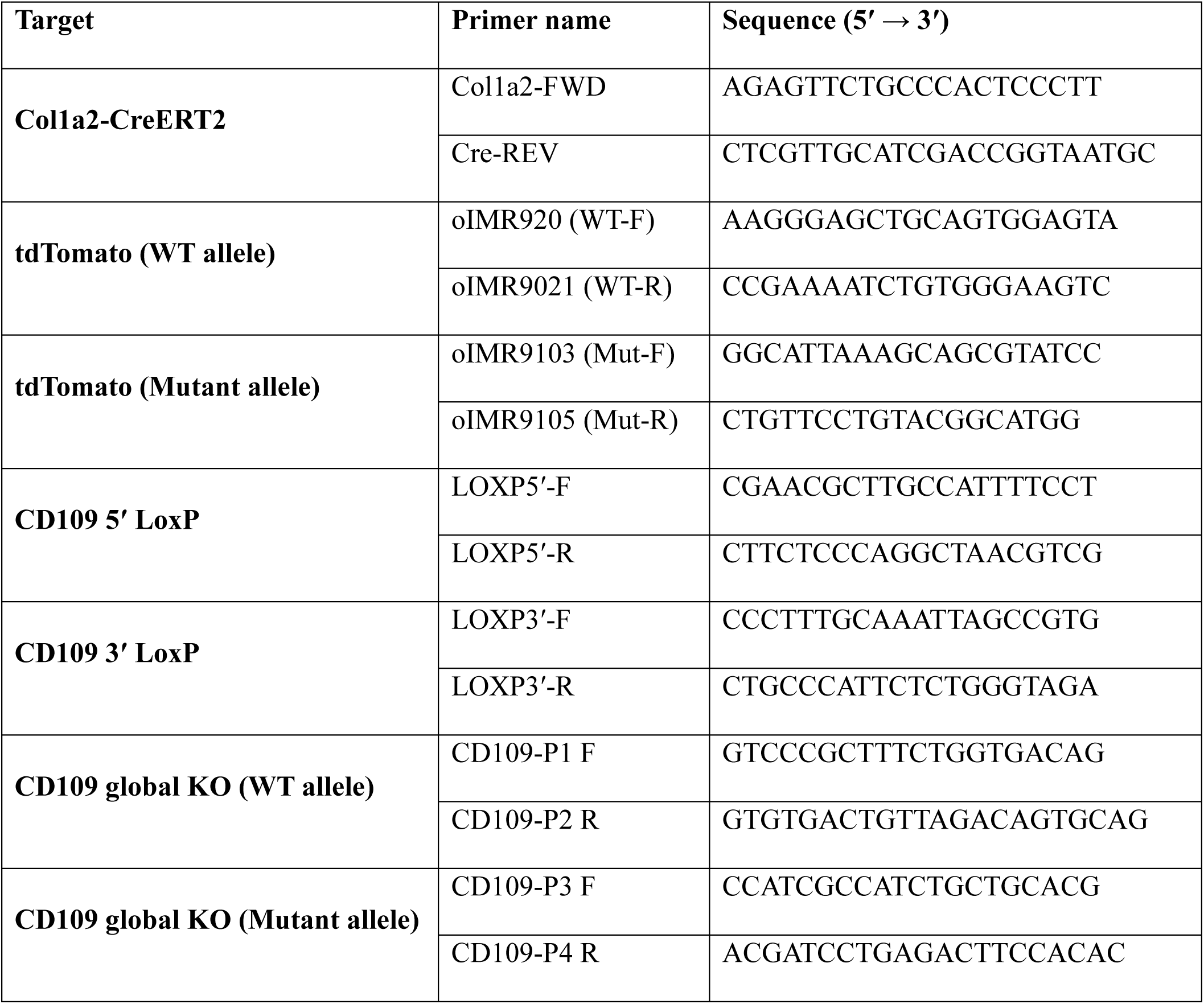
Primer sequences used for mouse genotyping.

### Histology and Immunohistochemistry (IHC)

Skin tissues were harvested, fixed, and processed for histological analysis. 5 µm sections were cut and mounted on glass slides for immunohistochemical staining. Sections were deparaffinized in xylene and rehydrated through graded ethanol solutions, followed by antigen retrieval using heat-induced epitope retrieval. Endogenous peroxidase activity and nonspecific binding were blocked prior to antibody incubation. Tissue sections were incubated with primary antibodies against F4/80 (macrophages), Ly-6G (neutrophils), CD3 (T cells), BST2 (plasmacytoid dendritic cells; pDCs), and IκBα (**Table 2**). Following primary antibody incubation, sections were incubated with appropriate horseradish peroxidase (HRP)-conjugated secondary antibodies. Signal detection was performed using 3,3′-diaminobenzidine (DAB) substrate and chromogen. Sections were subsequently counterstained with hematoxylin, dehydrated, and mounted. Stained sections were imaged using bright-field microscopy. Quantitative analysis of immunohistochemical staining was performed using Fiji (ImageJ). For all analyses, only the dermal compartment of skin sections was quantified. The epidermis and underlying adipose and muscle layers were manually masked and excluded from analysis to prevent their inclusion in threshold-based measurements.

**Table 2:** Primary and Secondary Antibodies used.

| **Antibody** | **Supplier** | **Host/Source** | **Technique and Dilution** |
| --- | --- | --- | --- |
| [KO-validated] NF-κB p65/RelA rabbit monoclonal antibody (A19653) | ABclonal | Rabbit | Western Blot (1:5000) |
| Phospho-NF-κB p65 (Ser276) rabbit polyclonal antibody (AP0123) | ABclonal | Rabbit | Western Blot (1:500) |
| IκBα rabbit monoclonal antibody (A19714) | ABclonal | Rabbit | Western Blot (1:1000)<br>IHC (1:100) |
| Phospho-IκBα-S32 Rabbit mAb (AP0707) | ABclonal | Rabbit | Western Blot (1:500) |
| TNFR1/TNFRSF1A rabbit polyclonal antibody (A1540) | ABclonal | Rabbit | Western Blot (1:500)<br>IP (2 μg/500 μg total protein)<br>IF (1:100) |
| TNFR2/TNFRSF1B rabbit monoclonal antibody (A19127) | ABclonal | Rabbit | Western Blot (1:1000)<br>IF (1:100) |
| TLR2 rabbit monoclonal antibody (A19125) | ABclonal | Rabbit | Western Blot (1:1000)<br>IF (1:100) |
| TLR4 rabbit polyclonal antibody (A5258) | ABclonal | Rabbit | Western Blot (1:1000)<br>IF (1:100) |
| BST2 rabbit monoclonal antibody (A23445) | ABclonal | Rabbit | IHC (1:100) |
| HRP-conjugated Goat anti-Rat IgG (H+L) (AS028) | ABclonal | Goat | IHC (1:500) |
| p-NFκB p50 Antibody (A-8): sc-271908 | Santa Cruz Biotechnology | Mouse | Western Blot (1:500) |
| NFκB p50 Antibody (E-10): sc-8414 | Santa Cruz Biotechnology | Mouse | Western Blot (1:500) |
| CD109 mouse monoclonal antibody (clone C-9; sc-271085) | Santa Cruz Biotechnology | Mouse | Western Blot (1:500)<br>IP (2 μg/500 μg total protein)<br>IF (1:100) |
| TNF-R2 Antibody (D-2): sc-8041 | Santa Cruz Biotechnology | Mouse | IP (2 μg/500 μg total protein) |
| Anti-Mouse F4/80 Purified (BM8, 14-480182) | eBioscience™ | Rat | IHC (1:100) |
| Anti-Mouse Ly-6G (Gr-1) Purified (RB6-8C5, 14-5931-85) | eBioscience™ | Rat | IHC (1:100) |
| Anti-Mouse CD3 Purified (17A2, 14-0032-81) | eBioscience™ | Rat | IHC (1:100) |
| CD109 (E4I2V) Rabbit Monoclonal Antibody #24765 | Cell Signaling | Rabbit | IHC (1:200) |
| Anti-rabbit IgG, HRP-linked antibody (#7074) | Cell Signaling | Goat | Western Blot (1:1000) |
| Anti-mouse IgG, HRP-linked antibody (#7076) | Cell Signaling | Horse | Western Blot (1:1000) |
| Rabbit IgG Horseradish Peroxidase-conjugated Antibody (HAF008) | R & D Systems | Goat | IHC (1:20) |
| Goat anti-Rabbit IgG (H+L), Alexa Fluor™ 594 | Invitrogen | Goat | IF (1:100) |
| Goat anti-Mouse IgG (H+L), Alexa Fluor™ 488 | Invitrogen | Goat | IF (1:100) |

### Cell Culture

Human hTERT-immortalized skin fibroblasts (0.2 × 10^6^ cells) were cultured in Dulbecco’s Modified Eagle Medium (DMEM) supplemented with serum and maintained under standard cell culture conditions. Cells were transfected with siRNA targeting CD109 or a non-targeting scrambled siRNA using Lipofectamine™ RNAiMAX Transfection Reagent. For transfection, cells were incubated in Opti-MEM for 4 hours in the presence of 7 nM Lipofectamine™ RNAiMAX, after which complete culture medium was restored. Following transfection, cells were serum-starved for 24 to 48 hours prior to treatment. For inflammatory signaling assays, cells were treated with tumor necrosis factor-α (TNF-α; 10 ng/mL) for 5 minutes, 10 minutes, or 60 minutes, depending on the downstream marker analyzed. Cells were harvested at the indicated time points for subsequent analyses.

### Protein Extraction and Quantification

Cells were lysed using Thermo Scientific™ RIPA Lysis and Extraction Buffer or Pierce™ IP Lysis Buffer according to the manufacturer’s instructions. Lysates were clarified by centrifugation, and supernatants were collected for downstream analyses. Total protein concentration was determined using the Quick Start™ Bradford 1× Dye Reagent (Bio-Rad, #5000205) following the manufacturer’s protocol. Absorbance was measured using a microplate reader, and protein concentrations were calculated using a bovine serum albumin (BSA) standard curve.

### SDS-PAGE and Western Blotting

Equal amounts of protein were resolved by SDS-PAGE alongside Precision Plus Protein™ Kaleidoscope™ Prestained Protein Standards (Bio-Rad). Proteins were subsequently transferred onto 0.45 µm nitrocellulose membranes (Amersham™ Protran™ Premium) at 90 V. Membranes were blocked in 1% BSA and incubated with primary antibodies diluted in BSA-containing blocking buffer (**Table 2**). Following primary antibody incubation, membranes were incubated with appropriate secondary antibodies (**Table 2**). The densitometry of immunoblots was measured using ImageJ software (NIH).

### Immunofluorescence

Cells were fixed using 4% paraformaldehyde (Electron Microscopy Sciences), washed with phosphate-buffered saline (PBS), and blocked in 1% BSA to reduce nonspecific binding. Samples were incubated with primary antibodies diluted in blocking buffer, followed by incubation with species-appropriate fluorescent secondary antibodies. Secondary antibodies used included Goat anti-Rabbit IgG (H+L), Alexa Fluor™ 594, and Goat anti-Mouse IgG (H+L), Alexa Fluor™ 488. After washing in PBS, samples were mounted using SlowFade™ Gold Antifade Mountant with DAPI for nuclear counterstaining and fluorescence preservation. Fluorescent images were acquired using a Zeiss LSM 780 Laser Scanning Confocal Microscope.

### Co-Immunoprecipitation

Cell lysates were prepared as described above and precleared with Protein G MagBeads (GenScript) to reduce nonspecific binding. Precleared lysates were then incubated with beads conjugated to a specific primary antibody or to an isotype-matched IgG control. Following incubation, immune complexes were captured using magnetic separation and washed to remove unbound proteins. Bound proteins were eluted from the beads and analyzed by SDS-PAGE and Western blotting as described above.

### Cytokine Array Analysis

CD109 knockdown was performed in fibroblasts as described above (Cell culture section). Conditioned medium was harvested, centrifuged at 3000 rpm for 10 min to remove debris, and cryopreserved at −80°C. To assess inflammation markers, the Human Inflammation Array Q3 (QAH-INF-3; RayBiotech, Inc.) was employed following standard protocols. Fluorescence scanning and subsequent data analysis were conducted using RayBiotech scanning services and Q-Analyzer software, respectively.

## Statistical Analysis

Statistical analyses were performed using GraphPad Prism (GraphPad Software, San Diego, CA, USA). Data were analyzed using either Student’s *t* test, Welch’s *t* test, or one-way analysis of variance (one-way ANOVA), as appropriate. The specific statistical test used for each experiment is indicated in the corresponding figure legends. Data are presented as mean ± standard error of the mean (SEM), and a *p* value < 0.05 was considered statistically significant.

## Availability of data and materials

Further information and requests for resources and reagents should be directed to and will be fulfilled by the corresponding author, Dr. Anie Philip. Single-cell datasets analyzed in this study are publicly available and were obtained from the Human Protein Atlas (https://www.proteinatlas.org) and the published pan-disease fibroblast atlas (https://celltype.info/project/613).

## Acknowledgments

We thank Mitra Cohen and team at the McGill Integrated Core for Animal Modeling (MICAM), Rosalind and Morris Goodman Cancer Institute at McGill University for the generation of CD109^fl/fl^ mice. We also thank members of the Philip laboratory for technical assistance and scientific discussions. We are grateful to Dr. Masahide (Nagoya University Graduate School of Medicine) for providing the global CD109 knockout mouse line. We also acknowledge the contributors of the Human Protein Atlas and the pan-disease fibroblast atlas for making their datasets publicly available. A.B. was supported by scholarships from the Canadian Institutes of Health Research (CIHR CGS-M), the Fonds de recherche du Québec – Santé (FRQS), and the Fonds de recherche du Québec – Nature et technologies (FRQNT). This work was supported by funding from the Canadian Institutes of Health Research (CIHR) Project Grant RN442412.

## Author contributions

Conceptualization: A.B., A.P.; Methodology: A.B., S.P., S.Z.; Investigation: A.B., S.P.; Formal Analysis: A.B.; Resources: A.P., GB; Writing – Original Draft: A.B.; Writing – Review & Editing: A.P., GB; Supervision: A.P.; Funding Acquisition: A.P.

## Declaration of generative AI and AI-assisted technologies

During the preparation of this manuscript, ChatGPT 5.2 was used solely to support language refinement and improve clarity of presentation. The graphical abstract was generated using *FigureLabs.ai*.

## Supplementary Figures

**Supplementary Figure 1:**
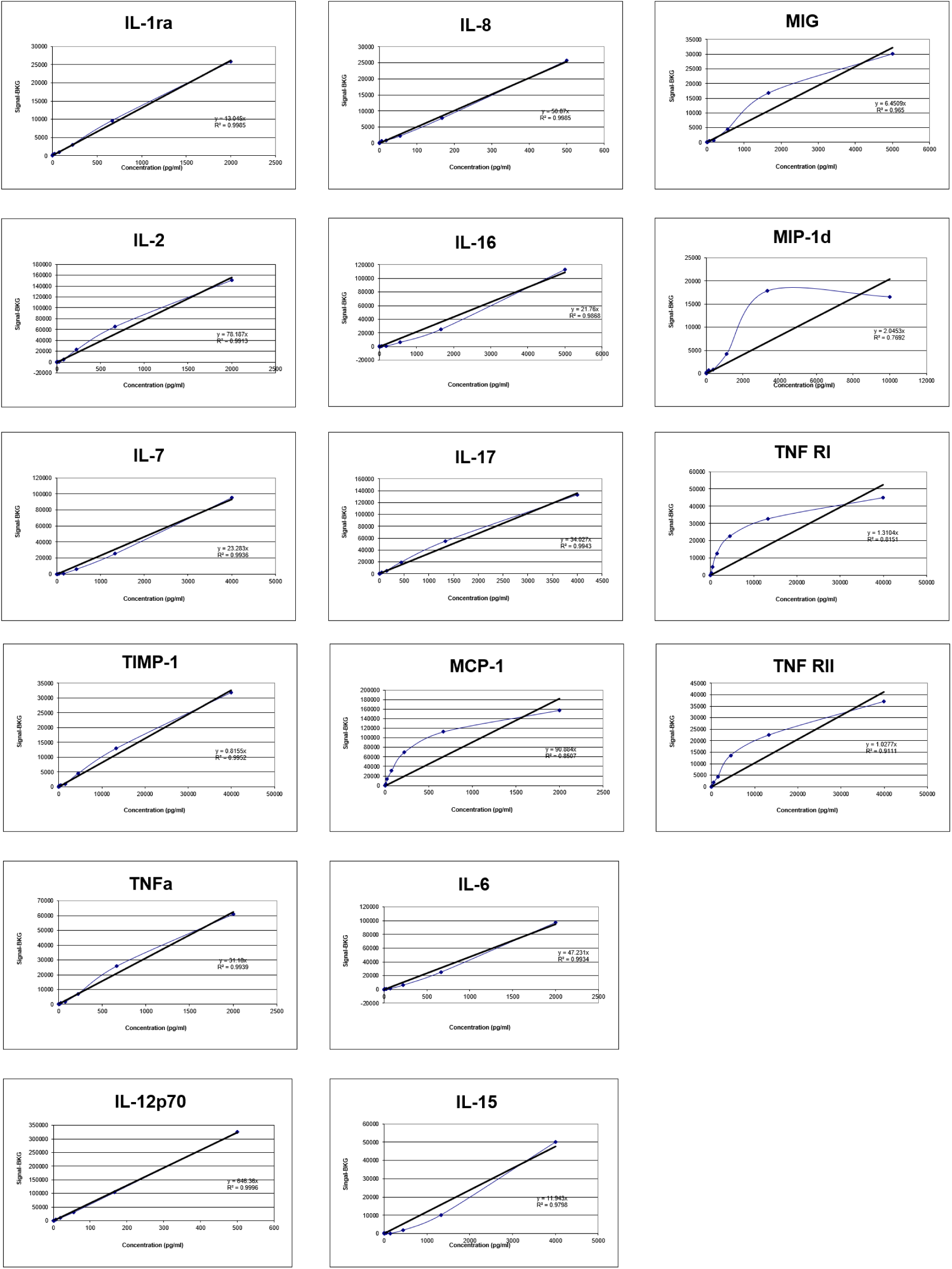
Standard curves for individual cytokines and soluble receptors. The cytokine levels were measured using the Quantibody® Human Inflammation Array 3 and the standard curves were generated using serial dilutions of recombinant standards provided by the manufacturer. Shown are representative calibration curves for IL-1 receptor antagonist (IL-1ra), IL-8, MIG (CXCL9), IL-2, IL-16, MIP-1δ, IL-7, IL-17, TNF receptor I (TNFRI), TIMP-1, MCP-1, TNF receptor II (TNFRII), TNF-α, IL-6, IL-12p70, and IL-15. Fluorescent signal intensity (y-axis) was plotted against analyte concentration (pg/mL; x-axis), and concentrations were calculated by linear regression within the validated dynamic range of each analyte. The fitted regression line and corresponding coefficient of determination (R²) are shown for each cytokine, indicating strong linearity across the working concentration range. These standard curves were used to calculate absolute cytokine concentrations in fibroblast-conditioned media following background subtraction and normalization.

**Supplementary Figure 2:**
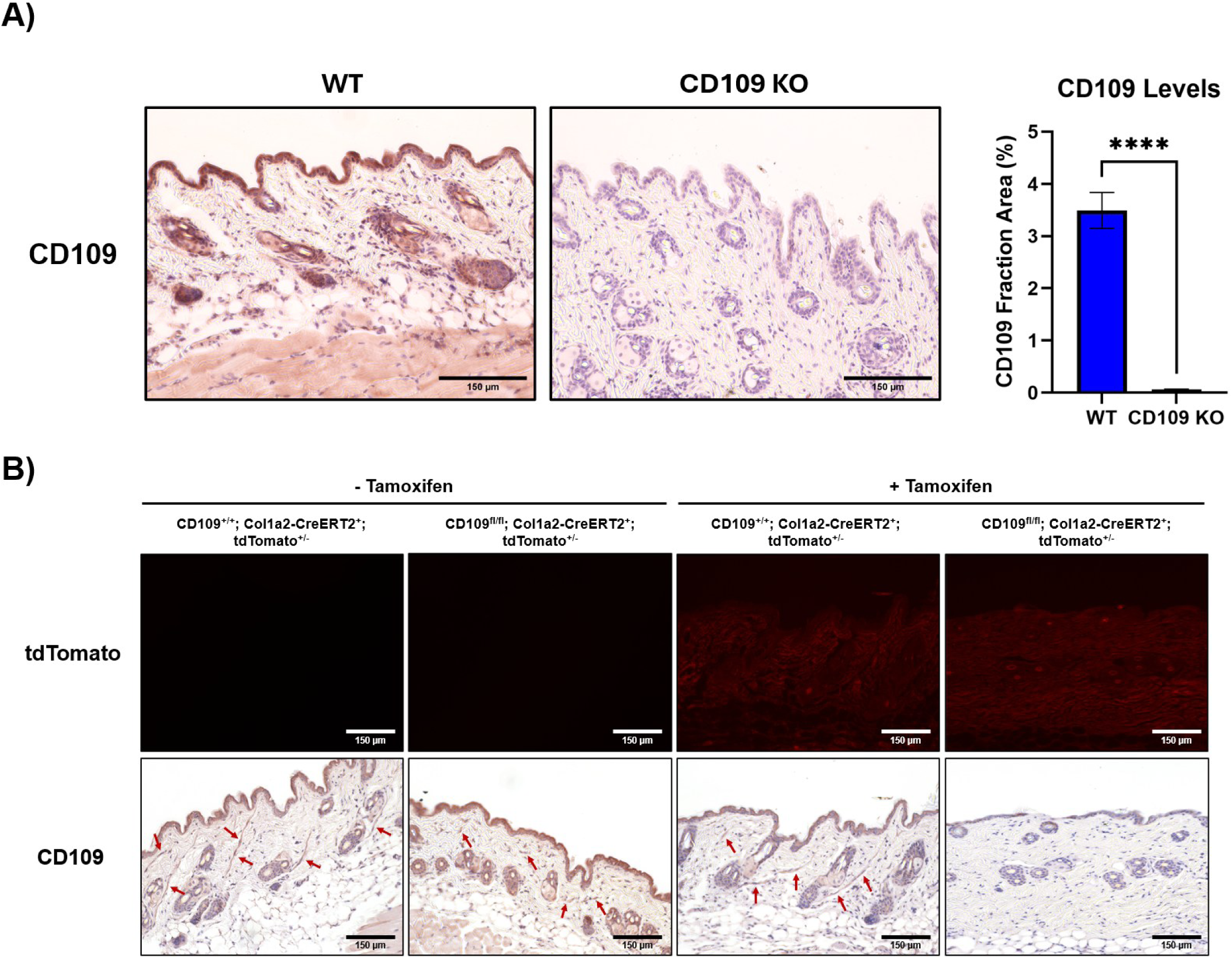
Confirmation of CD109 KO in mouse skin. **A)** Immunohistochemical staining for CD109 in WT and global CD109 KO mouse skin. Quantification of the CD109-positive area is shown to the right. **B)** Endogenous tdTomato expression in mouse skin. **C)** Immunohistochemical staining for CD109 in conditional CD109 KO mouse skin. Quantification of the CD109-positive area is shown to the right. Data: Mean ± SEM (n=3 per group). Statistical significance was assessed using one-way ANOVA (****p < 0.0001). Scale bars, 150 μm.

**Supplementary Figure 3:**
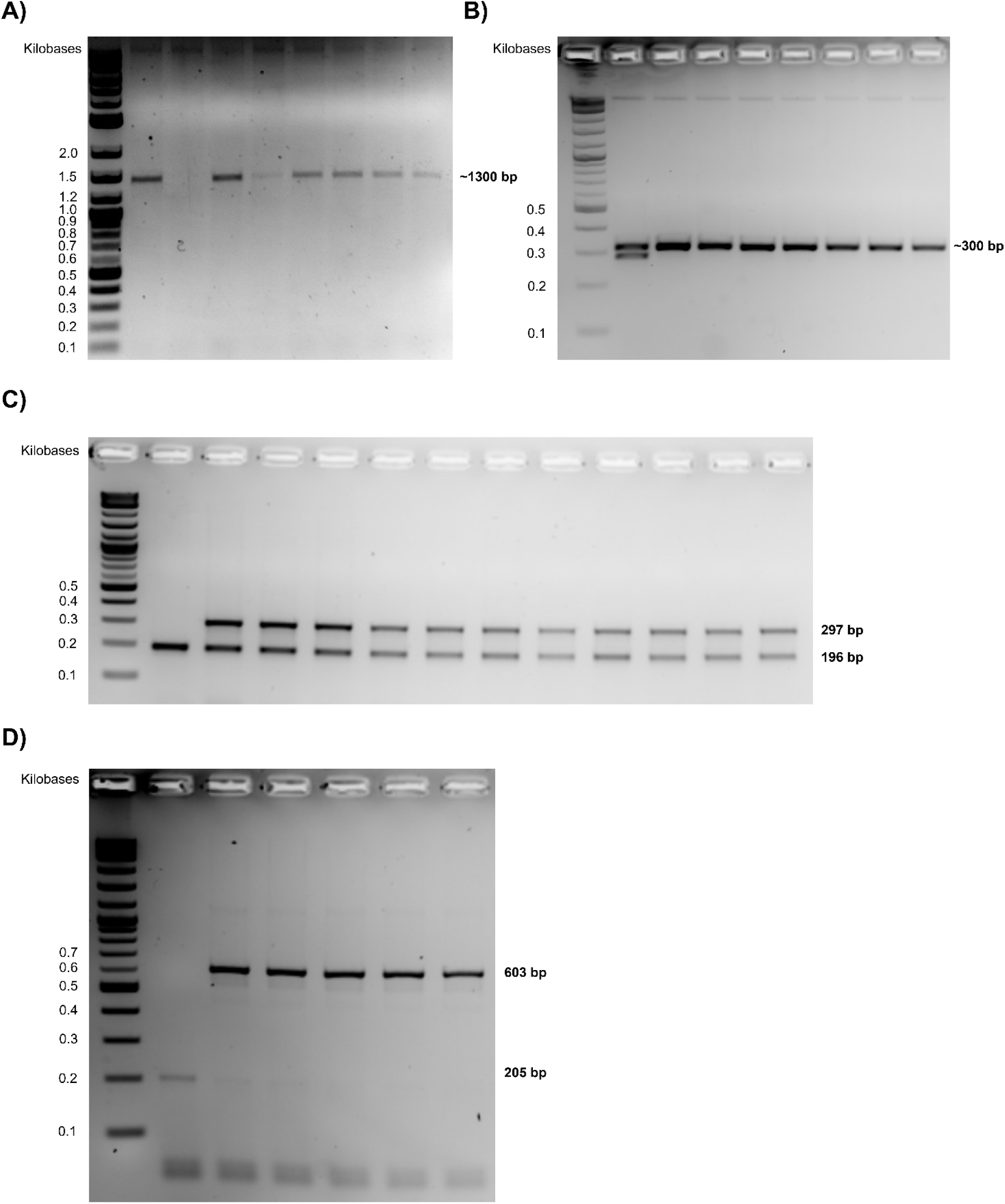
PCR-based genotyping of Col1a2-CreERT2, CD109 floxed alleles, tdTomato reporter, and global CD109 knockout mice. Representative agarose gel electrophoresis images showing PCR-based genotyping used to validate mouse lines employed in this study. **A)** Detection of the Col1a2-CreERT2 transgene using Cre-specific primers, yielding the expected Cre amplicon in Cre-positive animals. **B)** Verification of correct insertion of 5′ and 3′ loxP sites flanking the Cd109 locus using loxP-specific primer sets. Amplification of ∼300 bp fragments confirms the presence of the floxed Cd109 allele. **C)** Genotyping of the tdTomato reporter allele, showing PCR products corresponding to the wild-type (297 bp) and tdTomato knock-in (196 bp) alleles, allowing discrimination of WT, heterozygous, and homozygous reporter mice. **D)** Genotyping of the global CD109 KO allele, with PCR amplification yielding distinct products for the wild-type allele (205 bp) and the mutant KO allele (603 bp).

